# Time-varying mediation analysis for incomplete data with application to DNA methylation study for PTSD

**DOI:** 10.1101/2024.02.06.579228

**Authors:** Kecheng Wei, Fei Xue, Qi Xu, Yubai Yuan, Yuexia Zhang, Guoyou Qin, Agaz H. Wani, Allison E. Aiello, Derek E. Wildman, Monica Uddin, Annie Qu

## Abstract

DNA methylation (DNAm) has been shown to mediate causal effects from traumatic experiences to post-traumatic stress disorder (PTSD). However, the scientific question about whether the mediation effect changes over time remains unclear. In this paper, we develop time-varying structural equation models to identify cytosine-phosphate-guanine (CpG) sites where DNAm mediates the effect of trauma exposure on PTSD, and to capture dynamic changes in mediation effects. The proposed methodology is motivated by the Detroit Neighborhood Health Study (DNHS) with high-dimensional and longitudinal DNAm measurements. To handle the non-monotone missing DNAm in the dataset, we propose a novel Longitudinal Multiple Imputation (LMI) method utilizing dependency among repeated measurements, and employ the generalized method of moments to integrate the multiple imputations. Simulations confirm that the proposed method outperforms existing approaches in various longitudinal settings. In DNHS data analysis, our method identifies several CpG sites where DNAm exhibits dynamic mediation effects. Some of the corresponding genes have been shown to be associated with PTSD in the existing literature, and our findings on their time-varying effects could deepen the understanding of the mediation role of DNAm on the causal path from trauma exposure to PTSD risk.

## 1 Introduction

Post-traumatic stress disorder (PTSD) is a serious mental health disorder that arises after an individual witnesses or experiences traumatic events, such as severe accidents, natural disasters, or warfare. Although the causality of trauma on PTSD has been established from the biological perspective, the trauma-PTSD relation has not been elucidated on the population level of public health data. World Health Organization World Mental Health Surveys show that, although 69.7% of the surveyed population were exposed to at least one traumatic event, the cross-national lifetime prevalence of PTSD was only 5.6% among the trauma-exposed subpopulation (Koenen et al. 2017). This significant disparity of prevalence between trauma exposure and PTSD implies that there are potential intermediate factors intervening in the trauma-PTSD pathway to explain individual vulnerability to trauma and tendency to develop PTSD. On the other hand, extensive meta-analyses show that the variability in PTSD on the population level is not sufficiently explained by incorporating phenotype variables as predictors (Tortella-Feliu et al. 2019). These limitations in previous PTSD studies lead us to further explore genotype information, such as genetic and relevant molecular variation, as intermediate variables, and identify causal mediation pathways from trauma exposure to PTSD via the transmission of genetic variation.

DNA methylation (DNAm) is a molecular modification that alters gene expression without changing DNA sequence, and can be influenced by stress and socio-environmental factors (Turecki and Meaney 2016, Lussier et al. 2023). It has been found that aberrant DNAm is associated with a variety of stress-related neuropsychiatric disorders such as depression, schizophrenia, and PTSD (Wani et al. 2021, Wilker et al. 2023). Thus, DNAm may be a pathway by which traumatic events become biologically embedded and contribute to mental illness. Given the intermediate role of DNAm in mental disorder formulation, studies have suggested that DNAm may also mediate the trauma-PTSD mechanism: trauma increases or reduces DNAm levels, which in turn exacerbate or alleviate PTSD symptoms.

For example, Rutten et al. (2018) found that DNAm mediates the relationship between combat trauma and PTSD. Occean et al. (2022) showed that DNAm in the nuclear factor of activated T cells 1 mediates the exposure to different types of traumatic events and the risk for PTSD. Xue et al. (2022) identified the heterogeneous mediation effects of DNAm in different subpopulations.

From the perspective of mediation methodology, most of the existing mediation analyses on DNAm are cross-sectional based, i.e., only estimating mediation pathways at a single time point. However, DNAm levels and PTSD severity typically progress over time, therefore the relationships among trauma, DNAm, and PTSD may temporally change. For example, Wu et al. (2021) found that different pre-migration and post-migration stressors significantly affect refugees’ mental health at different resettlement stages. Lussier et al. (2023) demonstrated the temporal effects of adversity exposure on genetic variation across childhood and adolescence. Wilker et al. (2023) showed that the association between glucocorticoid receptor gene methylation status and PTSD symptoms is only significant in the early stages of study. Nevertheless, dynamic mediation analysis tools are still not well developed. By performing time-varying mediation analysis on longitudinal data, we can achieve a more comprehensive understanding of how causal pathways between variables in the disease process evolve over time: when the effects occur, diminish, strengthen, weaken, or change direction. Such insights could enrich understanding of the dynamic mediation effects of DNAm on pathways from trauma exposure to PTSD risk, and help to develop more targeted and time-sensitive interventions.

There have been some pioneering works about time-varying mediation analysis where exposure, mediator, and outcome are observed over multiple time points. For example, Rijnhart et al. (2022) and Cai et al. (2022) included time-interaction terms in their model to account for temporal effects. Zeng et al. (2021) and Zeng et al. (2022) extended the classical causal mediation method from a functional data analysis perspective. Ge et al. (2023) and Luo et al. (2023) studied dynamic mediation effects under a reinforcement learning framework. Nevertheless, most focus on cases where there is only one or a small number of mediators, with completely observed data. Few studies consider scenarios of high-dimensional mediators with incomplete data.

Missing data is ubiquitous in longitudinal study. In the statistical literature, two types of missingness structures are generally considered. One is called “monotone missingness” or “dropout”, where a subject may leave the study at a certain time point and never return. On the other hand, “non-monotone missingness” or “intermittent missingness” refers to when a subject may miss certain visits during follow-up and return at later visits. For our motivating Detroit Neighborhood Health Study (DNHS) which will be introduced later, subjects are supposed to have the methylation status of a large number of cytosine-phosphate-guanine (CpG) sites measured at each follow-up visit. Due to variability of compliance, the DNAm data exhibits the non-monotone missingness mechanism, and high-dimensional DNAm values are entirely missing at some visits by the individual. Classical approaches, such as weighting (Chen et al. 2021), imputation (Jahangiri et al. 2023), and likelihood-based methods (Tseng et al. 2016), have been developed to address nonmonotone missingness in longitudinal data. However, these approaches typically consider non-monotone missingness of the outcome or with a few covariates, and may not be directly applicable to our setting where high-dimensional DNAm values are entirely unobserved at some visits.

In this paper, we develop a new time-varying mediation method to investigate the temporal mediation effects of DNAm transmission from trauma to PTSD. We propose time-varying structural equation models, and adopt a regularization approach to select relevant mediators and identify time-varying effects, which yields several advantages. First, we can identify the important mediators from high-dimensional DNAm values at different time points, in contrast to existing dynamic mediation approaches considering only one or a small number of mediators (Rijnhart et al. 2022). Second, we can capture the trajectory of mediation effects even with a small number of time points, while existing approaches require dense time grids (Cai et al. 2022).

To handle non-monotone missing DNAm values, we propose a new imputation method called Longitudinal Multiple Imputation (LMI), which has the following major advantages. First, it makes full use of the dependence among repeated measurements, which exploits the inherent correlation structure of longitudinal data, not just based on low-rank structures or trajectory means (Mazumder, Hastie and Tibshirani 2010, Jahangiri et al. 2023). Second, it can handle high-dimensional mediators being entirely unobserved at some time points, in contrast to existing approaches which only consider missingness of the outcome or with a few covariates (Chen et al. 2021). Third, it does not require some individuals with complete repeated measurements in the dataset, whereas existing approaches may require that there must be some subjects with complete data, or that the data at the first time visit is always observed for all subjects (Jahangiri et al. 2023).

In simulation studies, the proposed method outperforms existing approaches under different missingness structures and different dimensions of mediators, and its performance is robust to different missingness mechanisms. In DNHS data analysis, our method identifies some DNAm CpG sites which exhibit dynamic mediation effects. Specifically, certain CpG sites initially have nonzero mediation effects, but the effects disappear over time. Certain CpG sites do not show mediation effects in early stages, but the effects emerge over time. Some genes (e.g., *PTPRK*) where selected CpG sites (e.g., cg09247979) are located have been reported as differentially expressed genes in previous PTSD meta-analyses (Chitrala, Nagarkatti and Nagarkatti 2016), and our further findings on their time-varying effects elucidate trauma-PTSD dynamic causal mechanisms.

The rest of the paper is organized as follows. Section 2 describes the background of the DNHS dataset. Section 3 introduces the proposed time-varying mediation models and the non-monotone missingness structure of mediators. Section 4 presents the proposed approaches for multiple imputation, data integration, and regularization. Section 5 illustrates the detailed implementation and algorithm. Section 6 includes the results of simulation studies. Section 7 applies the proposed method to the DNHS dataset. Section 8 concludes the research with some discussion.

## 2 Motivating application: Longitudinal trauma, PTSD, and DNAm

### 2.1 DNHS data

The Detroit Neighborhood Health Study (DNHS) is a prospective and representative longitudinal cohort study of predominantly African American adults living in Detroit, Michigan (Uddin et al. 2010, Goldmann et al. 2011). Participants completed a 40-minute structured telephone interview annually between 2008-2012 (i.e., waves 1-5), to assess demographic characteristics, exposure to traumatic events, and mental health status. All participants were given the opportunity to provide a specimen (venipuncture, blood spot, or saliva) for immune and inflammatory marker testing as well as genetic testing of DNA. We analyze the DNAm data with 526 subjects from waves 1, 2, 4, and 5 as there was no full blood draw in wave 3 as in the other waves. We refer to waves 1, 2, 4, and 5 as time points 1-4 throughout the paper.

The exposure variable of interest in our study is the number of traumatic events. In the initial interview (time point 1), participants were asked about their lifetime traumatic experiences using a predefined list of 19 traumatic events (Breslau et al. 1998). In subsequent interviews (time points 2-4), they were additionally queried about traumatic experiences since their last interview within the same list. At time points 2-4, we incorporate the number of traumatic events experienced since the previous interview to capture the cumulative severity of trauma exposure. The first plot in the left panel of Figure 1 shows the longitudinal trajectories of trauma from 15 subjects.

**Figure 1:**
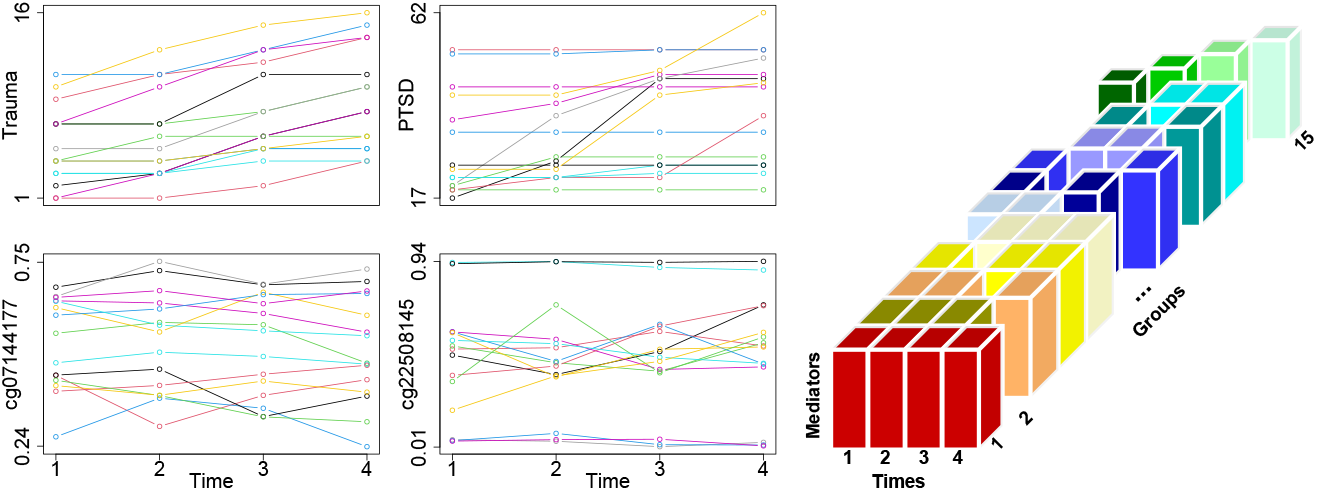
Left: Longitudinal trajectories of trauma, PTSD, and DNAm of two CpG sites from 15 subjects with complete repeated measurements of DNAm. Right: Non-monotone missingness structure of DNAm across all time points. Each blank column represents unobserved DNAm, while the colored ones represent observed DNAm.

The outcome variable of interest is PTSD symptom severity, which is measured through the widely used self-report PTSD Checklist (PCL, Ruggiero et al. 2003). The PCL contains 17 items corresponding to key symptoms of PTSD. In the initial interview (time point 1), participants were asked how much they had been bothered by each symptom using a 5-point scale (1-5) in reference to their worst lifetime traumatic experience, and the summary score for the 17 items were used to characterize the PTSD symptom severity. In subsequent interviews (time points 2-4), they were additionally queried about the worst traumatic experience since their last interview, and the summary score corresponding to it was similarly calculated. At time points 2-4, we choose the largest score calculated in previous and current interviews to capture the cumulative severity of PTSD symptoms. The second plot in the left panel of Figure 1 shows the longitudinal trajectories of PTSD from 15 subjects. We additionally incorporate demographic information such as age and gender, and blood work such as CD4 T cells and B cells, measured at baseline.

Existing studies suggest that epigenomic variations can be induced by trauma exposure and accompany the development of stress-related disorders (Rutten et al. 2018, Occean et al. 2022, Xue et al. 2022). However, few studies have reported brain-related epigenomic profiles that associate with PTSD risk. In DNHS, the methylation levels of 1879 CpG sites from paired blood and brain tissue (Braun et al. 2019) were assessed. We treat the DNAm levels of these 1879 CpG sites as potential mediators. The methylation level of a CpG site is quantified using the “Beta-value”, which ranges from 0 to 1 and is calculated as the proportion *M/*(*M* +*U* +100), where *M* and *U* represent the signal intensities of methylated and unmethylated probes (Mou et al. 2022). The third and fourth plots in the left panel of Figure 1 show the longitudinal trajectories of DNAm levels for two CpG sites from 15 subjects. We observe that the DNAm levels fluctuate within a specific range over the follow-up period, without exhibiting a clear trend or consistent pattern. The average rate of change in DNAm levels for all 1879 CpG sites across the 15 subjects between time points *t* and *t*^!^ can be calculated as 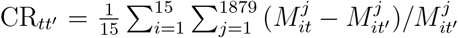. The corresponding results for CR_21_, CR_31_, CR_41_, CR_32_, CR_42_, and CR_43_ are 2.6%, 0%, 1.9%, −0.1%, 1.2%, and 3.2%, respectively.

The traumatic events and PTSD severity are easily assessed, so that we have complete repeated measurements for exposure and outcome. However, collecting and testing biospecimens is difficult and costly. Due to variability of compliance, loss to follow-up, and budget restriction, DNAm data could suffer from missingness. In DNHS, DNAm data of some subjects may be available only at certain visits, but are unobserved at other visits. The right panel of Figure 1 shows the non-monotone missingness structure of DNAm. Each blank column represents unobserved DNAm, while the colored ones represent observed DNAm. Participants can be divided into 15 groups, where group 1 has 4 repeated observations at all time points, while group 2 only has observations at time points 1-3, with high-dimensional DNAm data entirely missing at time point 4. The sample sizes of groups 1-15 are (15, 29, 7, 28, 21, 45, 27, 12, 23, 9, 21, 147, 24, 102, 16), respectively. There are 511 out of 526 subjects with missing values and the missingness rate of DNAm is 58.3%.

### 2.2 Novelty of our analysis

Several studies have suggested that DNAm plays a critical role as a mediator in the causal relationship between trauma and PTSD. For instance, Xue et al. (2022) leveraged DNHS data to identify multiple DNAm CpG sites mediating the effect of trauma on PTSD, highlighting that these epigenetic factors can play different mediation roles across different subpopulations. However, their analysis relied on baseline wave to estimate static mediation effects, which is not applicable for discovering potential time-varying genetic contributions to the disease process.

Time-varying effects are a common phenomenon in the genetic regulation of disease progression. Dynamic factors such as environmental changes and the natural course of the disease interact intricately with gene regulation mechanisms, suggesting that genetic variants may adapt to changing environmental conditions and show different effects at different stages of the disease (Lussier et al. 2023). Investigating time-varying genetic effects provides a more nuanced understanding of how genetic factors influence disease processes, helps identify critical time windows for intervention, and facilitates the design of time-sensitive therapeutic strategies.

However, studying time-varying effects poses great challenges. As illustrated in Figure 1, while trauma exposure and PTSD severity tend to increase monotonically over time, the dynamics of DNAm appear irregular, making it difficult to capture their patterns of variation. Moreover, the issues of missingness and high dimensionality within the DNHS dataset further complicate the analysis. These challenges underscore the necessity of developing innovative statistical models and estimation methods to characterize time-varying effects and accommodate complex data structures.

## 3 Models

In this section, we propose time-varying structural equation models to study dynamic mediation effects, and introduce the non-monotone missingness structure of mediators.

### 3.1 Time-varying structural equation models

Let 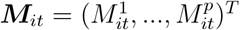 be a *p* × 1 vector of mediators, where 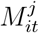 is the *j*-th (*j* = 1, …, *p*) mediator of the *i*-th (*i* = 1, …, *N*) subject, measured at the *t*-th (*t* = 1, …, *T*) time point.

Let *X*_*it*_ ∈ ℝ, *Y*_*it*_ ∈ ℝ, and ***Z***_*i*_ ∈ ℝ^*q*^ be the longitudinal exposure, the longitudinal outcome, and the demographic variables measured at baseline, respectively. We propose the following structural equation models:

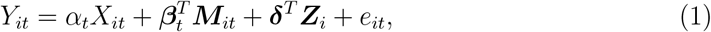

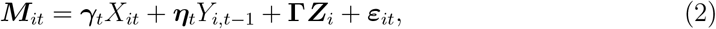

with *Y*_*i*0_ = 0, where the coefficients *α*_*t*_, 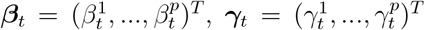, and 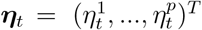 measure the time-varying effects at the *t*-th time point, ***δ*** ∈ ℝ^*q*^ and **Γ** ∈ ℝ^*p*×*q*^ measure the time-invariant effects. The random errors *e*_*it*_ and 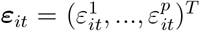 are zeromean, where *e*_*it*_ is independent of *X*_*it*_, ***M***_*it*_, and ***Z***_*i*_, and ***ε***_*it*_ is independent of *X*_*it*_, *Y*_*i,t*−1_, and ***Z***_*i*_. We define 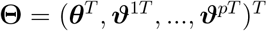 with 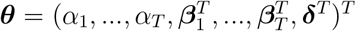 and 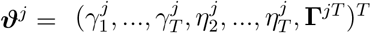, where **Γ**^*j*^ is the *j*-th row of **Γ** for *j* = 1, …, *p*.

Figure 2 illustrates the relationships between variables in models (1) and (2) without presenting ***Z***_*i*_, where solid unidirectional arrows represent causal relationships, while dashed bidirectional arrows indicate correlations. For a given time point *t, α*_*t*_ represents the direct effect from the exposure *X*_*it*_ to the outcome *Y*_*it*_, and 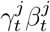 represents the indirect effect through the mediator 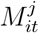. There are correlations among the mediators 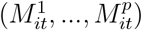. In the context of our real data, the DNAm levels of multiple CpG sites exhibit correlations, reflecting their shared regulatory roles or proximity to similar biological functions (Mou et al. 2022).

**Figure 2:**
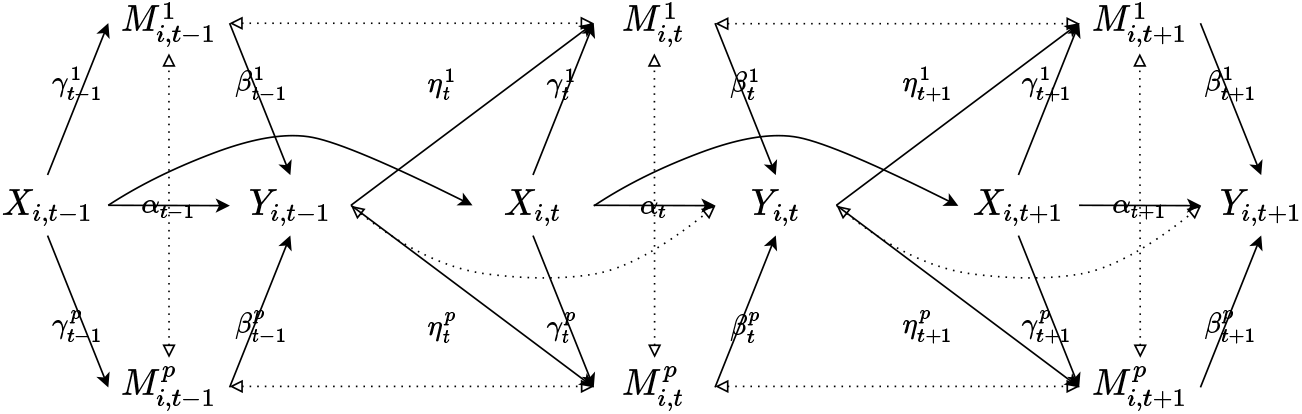
Dynamic mediation structure among time points *t* − 1, *t*, and *t* + 1. Solid unidirectional arrows represent causal relationships, while dashed bidirectional arrows indicate correlations.

For adjacent time points *t* − 1 and *t*, we assume both correlation and causation exist among the exposure, mediators, and outcome. We postulate correlations among repeated measurements of outcome *Y*_*it*_. And similarly, we assume correlations among repeated measurements of mediator 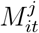. Regarding the causal relationships across time points, since *X*_*it*_ reflects the accumulated level of trauma exposure, it is influenced mainly by the exposure at the previous time point. Based on existing literature, DNAm is primarily influenced by external adverse environmental exposures or disease conditions (Turecki and Meaney 2016), we specify the causal effect of PTSD condition *Y*_*i,t*−1_ on 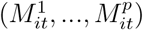 to be 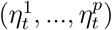. Following the Markov assumption (Luo et al. 2023), we assmue that spillover effects among variables occur exclusively at adjacent time points.

Regression coefficients in models (1) and (2) can be linked to causal estimands. Let *Y* ^*x*,***m***^ be the potential outcome that would have been observed if *X* was set to *x* and ***M*** was set to ***m***, and ***M*** ^*x*^ be the potential mediators that would have been observed if *X* was set to *x*. Following Bind et al. (2016), we formalize the following assumptions:

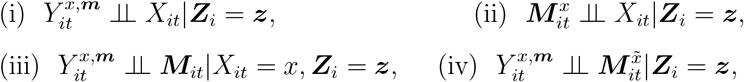

which indicate that there is no (i) unmeasured exposure-outcome confounding, (ii) exposure-mediator confounding, (iii) mediator-outcome confounding, or (iv) exposureinduced confounding, where 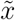 is the realization of exposure at a different value from *x*. Corresponding to our DNHS data, to estimate the direct effect of trauma exposure *X*_*it*_ on PTSD outcome *Y*_*it*_, as well as the indirect effect mediated through DNAm ***M***_*it*_, it is essential to control for confounding bias. Assumption (i) requires that all causes of trauma and PTSD are observed and adjusted for. These causes may include demographic information such as age and gender (Wani et al. 2021), which have been accounted for in ***Z***_*i*_. Assumption (ii) requires that all causes of trauma and DNAm are observed and adjusted for. These may include demographic information like age and gender (Wani et al. 2021), which have been accounted for in ***Z***_*i*_. Assumption (iii) requires that all causes of DNAm and PTSD are observed and adjusted for. These may include demographic information such as age and gender, as well as blood work (Occean et al. 2022), which have been accounted for in ***Z***_*i*_. Assumption (iv) requires that the causes (except for trauma) of DNAm and PTSD are not induced by trauma. This assumption is reasonable, as the demographic information and blood work we control for are intrinsic attributes of individuals. Note that although *Y*_*i,t*−1_ may have a spillover effect on 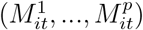, it does not directly affect *X*_*it*_ and *Y*_*it*_.

Consequently, there is no time-varying confounding in our study, similar to the arguments by VanderWeele and Tchetgen Tchetgen (2017), and thus following Bind et al. (2016), the natural direct effect at time point *t* for *x* versus 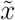 is 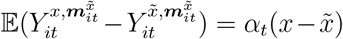, and the natural indirect effect at time point *t* for *x* versus 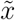 is 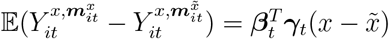.

### 3.2 Non-monotone missing mediators

In our DNHS data, exposure *X*_*it*_, outcome *Y*_*it*_, and confounders ***Z***_*i*_ are always observed, but the mediators ***M***_*it*_ suffer from non-monotone missingness due to the following reasons: ***M***_*i*_ = (***M***_*i*1_, …, ***M***_*iT*_)^*T*^ of some subjects may be available only at certain visits, but missing at the next time point, and measured again at later visits. Given the non-monotone missingness, subjects can be divided into *R* disjoint groups based on different missingness patterns across all time points. Figure 3 illustrates an example where we have *T* = 3 time points and *R* = 7 groups. Each blank column represents unobserved mediators, while the colored ones represent observed mediators. Notice that subjects in group 1 have observed ***M***_*i*_ at all time points, but subjects in group 2 have observed ***M***_*i*_ only at time points 1 and 2.

**Figure 3:**
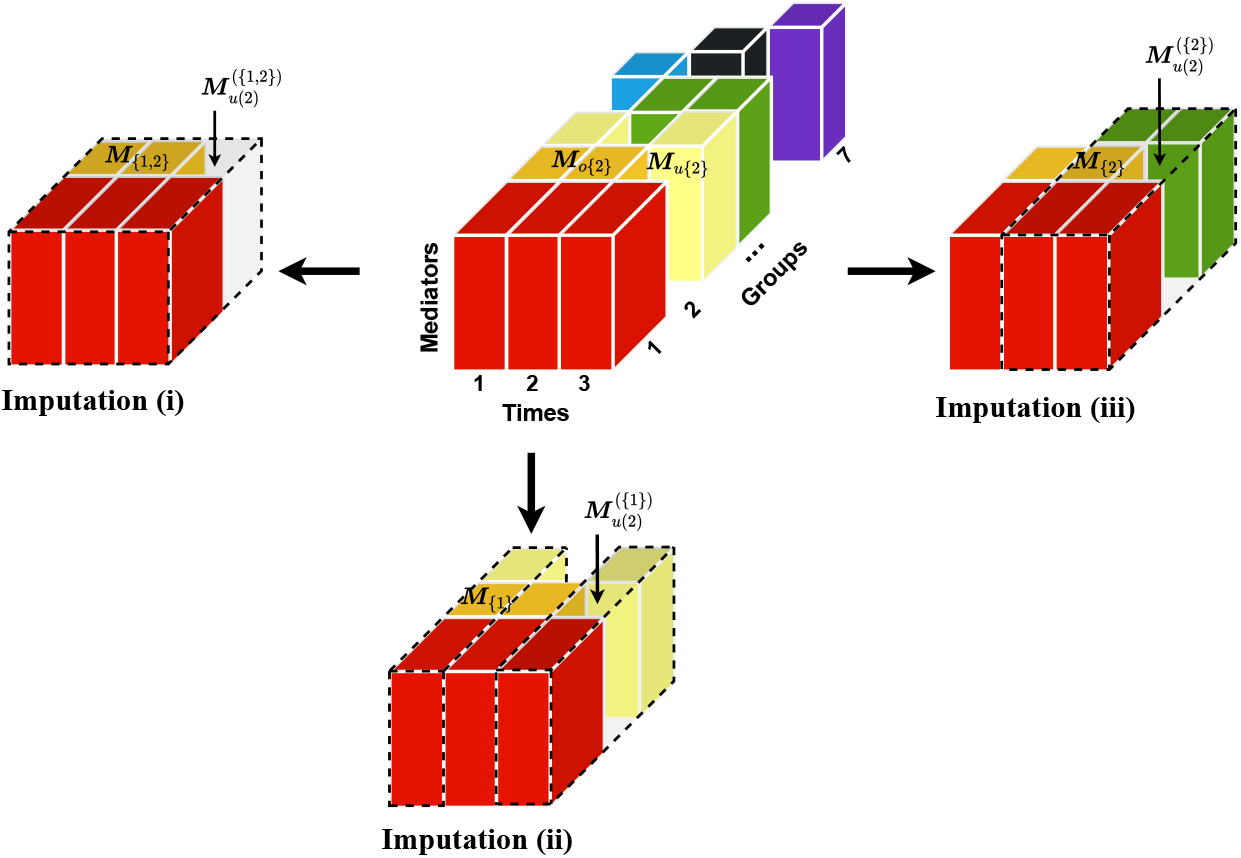
Middle: Non-monotone missingness structure of mediators across all time points. Each blank column represents unobserved mediators, while the colored ones represent observed mediators. Surroundings: Multiple imputations for the missing column ***M***_*u*(2)_ in group 2.

Define the missing indicator 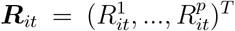, where 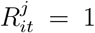 for *j* = 1, …, *p* if ***M***_*it*_ is observed at time point *t*, and 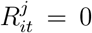 for *j* = 1, …, *p* if ***M***_*it*_ is missing at time point *t*. In this paper, we assume that the missingness mechanism is missing at random (MAR, Little and Rubin 2019), meaning that the conditional distribution of the missing indicator ***R***_*i*_ = (***R***_*i*1_, …, ***R***_*iT*_)^*T*^, denoted as *f*_***R***_ (***r***_*i*_|***X***_*i*_, ***Y***_*i*_, ***Z***_*i*_, ***M***_*i*_), depends only on the exposure ***X***_*i*_ = (*X*_*i*1_, …, *X*_*iT*_)^*T*^, the outcome ***Y***_*i*_ = (*Y*_*i*1_, …, *Y*_*iT*_)^*T*^, the confounders ***Z***_*i*_, and the observed mediator component ***R***_*i*_***M***_*i*_ (Wei et al. 2022).

Let ℋ(*r*) be the index set of subjects in group *r* and *n*_*r*_ = |ℋ(*r*)| be the sample size (*r* = 1, …, *R*). Let *o*(*r*) and *u*(*r*) be the index sets of time points corresponding to observed and unobserved mediators in group *r*. Consequently, ***M***_*o*(*r*)_ and ***M***_*u*(*r*)_ represent observed and unobserved mediators in group *r*. In addition, for each *t* ∈ *u*(*r*), we define ℐ(*r, t*) as the collection of time sets, and each element in ℐ(*r, t*) is a subset of *o*(*r*). The ℐ(*r, t*) can be obtained by the following steps: (i) let 𝒢(*r, t*) be the index set of the groups where mediators are observed not only at time point *t*, but also at least one time point in *o*(*r*);

(ii) for each *r*^!^ ∈ 𝒢(*r, t*), let 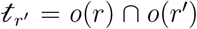; (iii) 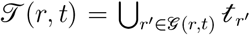. Note that the definition of ℐ(*r, t*) is to prepare for the imputation method later. That is, we can use observed values at each time set *t* ∈ ℐ(*r, t*) to impute the unobserved values at time point *t* ∈ *u*(*r*).

## 4 Methods

In this section, we first propose a new imputation method called Longitudinal Multiple Imputation (LMI) to handle non-monotone missing data. Second, we adopt the generalized method of moments (GMM, Hansen 1982) to combine the multiple imputations. Third, we propose a regularization approach to identify relevant mediators and estimate time-varying effects. Finally, we employ principal component analysis to solve the singularity issue of the weighting matrix in GMM.

### 4.1 Longitudinal multiple imputation (LMI)

In this subsection, we propose the LMI approach to handle non-monotone missing longitudinal data. The main idea is that, due to the correlation between repeated measurements ***M***_*t*_ (*t* = 1, …, *T*) from the same subject, we can use the observed values at some time points to predict the unobserved values at other time points. For example, for subjects with observed ***M***_*t*_ at time points 1 and 2, we can use some statistical methods, such as regression, to learn the correlation of variables between these two time points. Thus, for some other individuals with observed ***M***_*t*_ at time point 1 but missing ***M***_*t*_ at time point 2, we can impute the data at time point 2 based on the learned correlation.

Given a group *r* with a time point *t* ∈ *u*(*r*), subjects have unobserved *M*^*j*^ at this time point, but have observed ***M***_*t*_ at each time set *t* ∈ ℐ(*r, t*). We can use observed values at each time set *t* ∈ ℐ(*r, t*) to impute the unobserved values at time point *t* ∈ *u*(*r*). Specifically, given a time set *t* ∈ ℐ(*r, t*), we first estimate 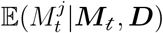 using other groups containing observed 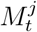 and ***M***_*t*_, where ***D*** contains exposure ***X*** = (*X*_1_, …, *X*_*T*_)^*T*^ at all time points, outcome *Y*_*t*−1_ at time point *t* − 1, and confounders ***Z***. Based on the learned correlation, we then impute unobserved *M*^*j*^ using ***M***_*t*_ and ***D***. Let 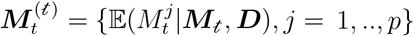 represent the imputed values based on the time set *t* ∈ ℐ(*r, t*), then we can obtain |ℐ(*r, t*)| imputations.

We illustrate the LMI with an example in Figure 3. Specifically, ℋ(2) refers to the index set of subjects in group 2. Since subjects in group 2 have observed mediators at time points 1 and 2, but have unobserved mediators at time point 3, *o*(2) = {1, 2} and ***M***_*o*(2)_ refers to observed mediators, *u*(2) = {3} and ***M***_*u*(2)_ refers to unobserved mediators.

The collection of time sets ℐ(2, 3) = {{1, 2}, {1}, {2}} since: (i) 𝒢(2, 3) = {1, 3, 4}; (ii) *t*_1_ = *o*(2) ∩ *o*(1) = {1, 2}, *t*_3_ = *o*(2) ∩ *o*(3) = {1}, and *t*_4_ = *o*(2) ∩ *o*(4) = {2}; (iii) 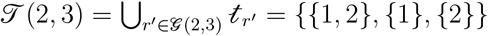.

Given group 2 with unobserved ***M***_*t*_ at time point *t* ∈ *u*(*r*) = 3, we can impute the unobserved values in |ℐ(*r, t*)| = |{{1, 2}, {1}, {2}}| = 3 ways: for *j* = 1, ‥, *p*, (i) for *t* = {1, 2}, we estimate 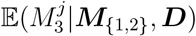 using group 1, and then obtain the imputed values 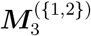(ii) for *t* = {1}, we estimate 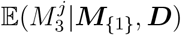 using groups 1 and 3, and then obtain the imputed values 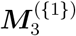;(iii) for *t* = {2}, we estimate 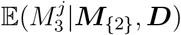 using groups 1 and 4, and then obtain the imputed values 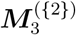.

#### Remark 1.

One novelty of our LMI method is to make full use of the dependence among repeated measurements, which exploits the inherent correlation structure of longitudinal data, not based on stringent low-rank assumptions (Mazumder, Hastie and Tibshirani 2010) or information-poor trajectory means (Jahangiri et al. 2023). The LMI method allows high-dimensional variables to be entirely unobserved at some time points, while existing approaches only consider missingness of the outcome or with a few covariates (Chen et al. 2021). Furthermore, the LMI method allows for the absence of individuals with complete repeated measurements in the dataset, whereas existing approaches may require that there must be some subjects with complete data, or that the data at the first time visit is always observed for all subjects (Jahangiri et al. 2023).

### 4.2 Combining multiple imputations

In this subsection, we propose to combine the information from all LMIs. Since different imputations correspond to different estimating functions, based on moment conditions (Hansen 1982), we integrate all information through the GMM.

First, we construct the moment conditions corresponding to model (1). For a given group *r, j* = 1, ‥, *p*, and each time set *t* ∈ ℐ(*r, t*), since unobserved *M*^*j*^ at time point *t* ∈ *u*(*r*) is imputed through 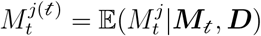, the covariates ***M***_*t*_ and ***D*** are uncorrelated with residuals of the projection 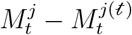. Therefore,

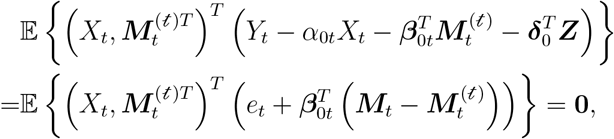

and

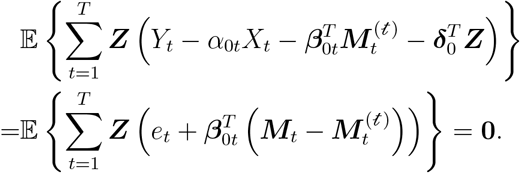

The moment conditions are satisfied for the true parameter 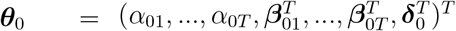.

We then construct estimating functions based on these moment conditions. For each *i* ∈ ℋ(*r*), let 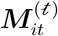 be the imputed values based on time set *t* ∈ ℐ(*r, t*), where 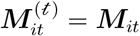 for *t* ∈ *o*(*r*), and 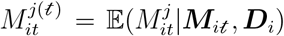 for *t* ∈ *u*(*r*) and *j* = 1, ‥, *p*. The estimating functions for *X*_*it*_ and 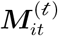 at time point *t* ∈ *o*(*r*) are 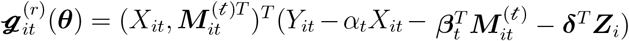 in which 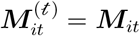 and the estimating functions for *X*_*it*_ and 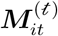 at time point *t* ∈ *u*(*r*) are 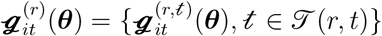, where

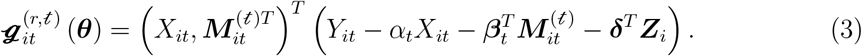

The estimating functions for ***Z***_*i*_ are 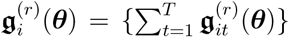, where 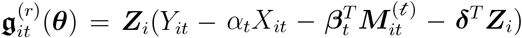 at time point *t* ∈ *o*(*r*) in which 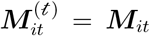, and 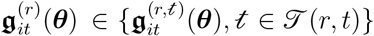 at time point *t* ∈ *u*(*r*) in which

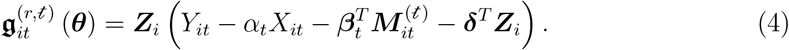

To integrate information from all groups, we propose an aggregated vector of estimating functions:

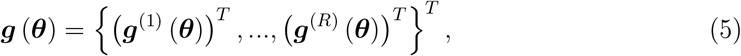

where 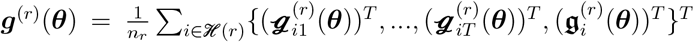. Note that the total number of estimating functions exceeds the dimension of parameters, and the estimating functions from groups with fewer missingness or more precise imputations tend to have smaller variance. To combine all estimating functions in ***g***(***θ***) and put more weights on high-quality moments, we propose the following quadratic loss function:

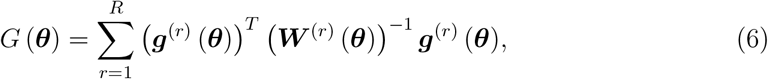

with weighting matrix 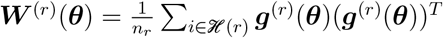.

For each mediator *M*^*j*^ (*j* = 1, …, *p*) in model (2), we can get the aggregated vector of estimating functions ***h***^*j*^(***ϑ***^*j*^) similar to (5). Detailed formulations are shown in the Supplementary Material. We propose the following quadratic loss function:

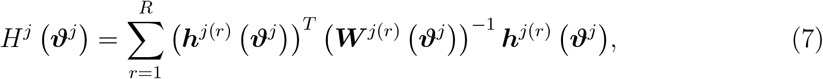

with weighting matrix 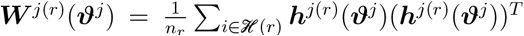. The loss function incorporating models (1) and (2) for all subjects is 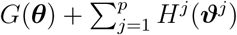.

#### Remark 2

For multiple imputed data, different imputations correspond to different estimating functions, all of which satisfy the moment conditions. We use GMM to combine all estimating functions, which adaptively puts more weights to imputations with smaller variance, while still consider the information contained in imputations with larger variance. In contrast, if we only use the imputation with the smallest variance while discarding the others, it may lead to efficiency losses due to not use of available information.

### 4.3 Regularization

To select the relevant mediators from high-dimensional variables and to estimate the timevarying effects, we adopt the regularization approach. Specifically, we propose to incorporate three penalty terms to: (i) identify mediators with large mediation effects at each time point, (ii) shrink similar effects at adjacent time points, and (iii) shrink effects of demographic variables. Consequently, the objective function of the proposed method is

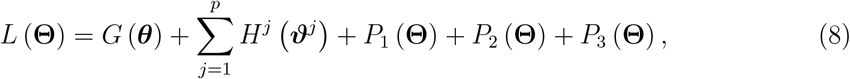

where

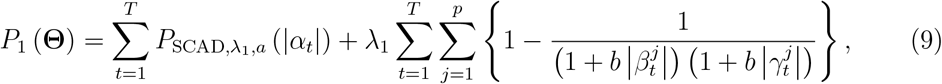

is a penalty for direct effects and indirect effects, with 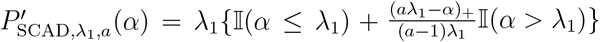 for some *α >* 0, *a >* 2, and *b >* 0,

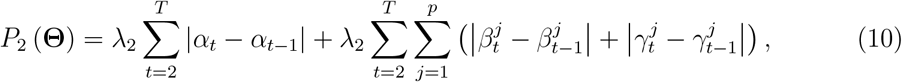

is a fusion penalty for time-varying effects, and

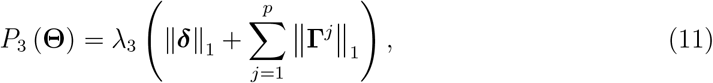

is a penalty for demographic effects. The regularization parameters *λ*_1_, *λ*_2_, and *λ*_3_ control the amount of shrinkage.

The second term in *P*_1_(**Θ**) is a bi-level mediation penalty for indirect effects, which links models (1) and (2) by penalizing ***β***_*t*_ and ***γ***_*t*_ jointly rather than separately, and achieves bilevel variable selection (Huang et al. 2009). This estimator not only obtains the sparse solution for 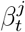 and 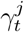 when 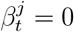 and 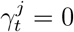, but also obtains the sparse solution for 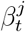 or 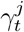 when 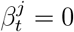 or 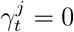. On the other hand, it inherits the advantage of the smoothly clipped absolute deviation (SCAD, Fan and Li 2001) penalty where the function gradually levels off as 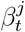 and 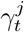 increase, which avoids over-shrinkage. In general, it is possible that not every mediator has different effects at different time points. We use the fused lasso penalty (Tibshirani et al. 2005) in *P*_2_(**Θ**) to shrink similar effects at adjacent time points, which links ***β***_*t*_ and ***γ***_*t*_ across all time points. *P*_3_(**Θ**) is a lasso penalty (Friedman, Hastie and Tibshirani 2010) for selecting demographic effects to avoid over-fitting.

### 4.4 Principal components of weighting matrix

The GMM estimators in (6) and (7) may have poor finite sample performance in highly overidentified models, due to imprecise estimation of the weighting matrix ***W*** ^(*r*)^(***θ***) or ***W*** ^*j*(*r*)^(***ϑ***^*j*^), or under the singularity of the weighting matrix (Doran and Schmidt 2006). This could be attributed to the large number of estimating functions compared to the relatively small sample size, or due to the overlap of information between LMIs. For example, as illustrated in Figure 3, the observed values at time point 1 in group 2 are used for the estimation of both 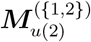 and 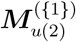 and the observed values at time point 2 in group 2 are used for the estimation of both 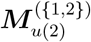 and 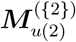.

To address the singularity issue, we reduce the dimension of ***g***^(*r*)^ and ***h***^*j*(*r*)^ (*r* = 1, …, *R* and *j* = 1, …, *p*), by combining informative estimating functions, for example, using the first several largest principal components. If 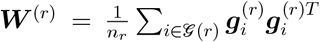 is singular or close to singular, we extract the first *u*^(*r*)^ principal components 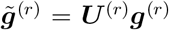 from ***g***^(*r*)^, where ***U*** ^(*r*)^ contains *u*^(*r*)^ eigenvectors of ***W*** ^(*r*)^ corresponding to the largest *u*^(*r*)^ nonzero eigenvalues. Similarly, if 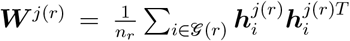 is singular or close to singular, we extract the first *u*^*j*(*r*)^ principal components 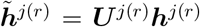 from ***h***^*j*(*r*)^, where ***U*** ^*j*(*r*)^ contains *u*^*j*(*r*)^ eigenvectors of ***W*** ^*j*(*r*)^ corresponding to the largest *u*^*j*(*r*)^ nonzero eigenvalues. The numbers of principal components *u*^(*r*)^ and *u*^*j*(*r*)^ (*r* = 1, …, *R* and *j* = 1, …, *p*) can be selected based on the information criterion proposed by Cho and Qu (2015) to capture sufficient information from the estimating functions. Note that if ***W*** ^(*r*)^ or ***W*** ^*j*(*r*)^ is not singular, ***U*** ^(*r*)^ or ***U*** ^*j*(*r*)^ is an identity matrix. Consequently, the final objective function is

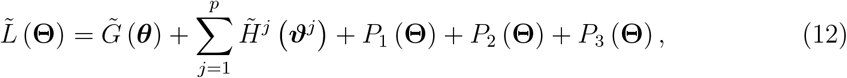

where the quadratic form 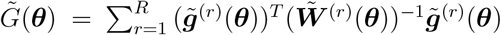 with transformed weighting matrix 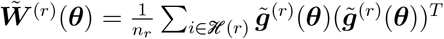, and the quadratic form 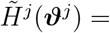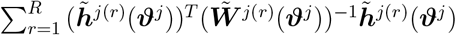 with transformed weighting matrix 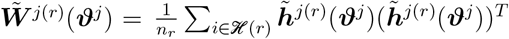.

## 5 Computation

In this section, we provide the computational details of the proposed method. We first describe the implementation of LMI from Section 4.1. Then we introduce an information criterion to select the numbers of principal components from Section 4.4. Next, we present a smoothing technique to handle the non-separable fusion penalty from Section 4.3. Afterwards, we introduce the overall optimization algorithm of the proposed method for a given set of regularization parameters (*λ*_1_, *λ*_2_, *λ*_3_)^*T*^ from Section 4.3. Finally, we propose an information criterion to select these regularization parameters.

When conducting LMI, the conditional expectation 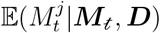 is required to be estimated for each time set *t* ∈ ℐ(*r, t*), time point *t* ∈ *u*(*r*), group *r* = 1, …, *R*, and dimension *j* = 1, …, *p*. In this paper, we adopt the lasso-regularized linear model (Friedman, Hastie and Tibshirani 2010) to account for high-dimensionality, where the regularization parameter is selected by cross-validation.

To capture most of the information from the estimating functions ***g***^(*r*)^ and ***h***^*j*(*r*)^ (*r* = 1, …, *R* and *j* = 1, …, *p*), we select the numbers of principal components *u*^(*r*)^ and *u*^*j*(*r*)^, by minimizing the following Bayesian-type information criterion (Cho and Qu 2015):

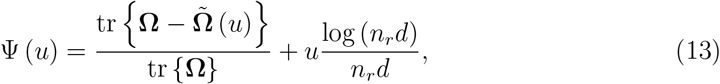

where tr{·} is the trace of a matrix, **Ω** represents the weighting matrix ***W*** ^(*r*)^ or ***W*** ^*j*(*r*)^ with dimension 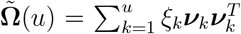 is an approximation of **Ω** based on the largest *u* eigenvectors, and *ξ*_*k*_ is the *k*-th largest eigenvalue of **Ω** corresponding to the eigenvector ***?***_*k*_. Since 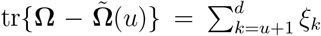, the minimizer of Ψ(*u*) is indeed the number of eigenvalues greater than tr{**Ω**}log(*n*_*r*_*d*)*/*(*n*_*r*_*d*).

Since the non-smooth and non-separable of fusion penalty *P*_2_(**Θ**) in (10) makes the optimization challenging, we use the smoothing technique from Chen et al. (2012). Specifically, we reformulate the fused lasso as 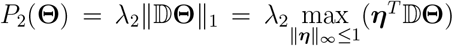, where 𝔻 is a difference operator corresponding to the differences in *P*_2_(**Θ**). Let 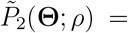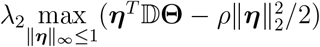, where *ρ* is a positive smoothing parameter and 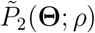 proximates *P*_2_(**Θ**) as *ρ* → 0. Define a projection operator *S*(*x*) = − 𝕀 (*x <* −1) + *x 𝕀* (−1 ≤ *x* ≤ 1) + 𝕀 (*x >* 1), we can deduce that ***η***^*^ = *S*(𝔻 **Θ***/ρ*) is indeed the optimal solution in 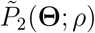. Thus we have

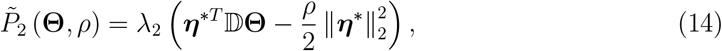

which is convex and differentiable in **Θ**. We set *ρ* = 10^−4^ following Chen et al. (2012). We propose an algorithm based on the iterated GMM with principal component analysis. Specifically, given **Θ**^(*s*−1)^ from the (*s* − 1)-th iteration, for *r* = 1, …, *R* and *j* = 1, …, *p*, if the weighting matrix ***W*** ^(*r*)^(***θ***^(*s*−1)^) or ***W*** ^*j*(*r*)^(***ϑ***^*j*(*s*−1)^) is singular or close to singular, we first select the number of principal components *u*^(*r*)(*s*−1)^ and *u*^*j*(*r*)(*s*−1)^ by minimizing the information criterion in (13), then construct principal component matrices ***U*** ^(*r*)^(***θ***^(*s*−1)^) and ***U*** ^*j*(*r*)^(***ϑ***^*j*(*s*−1)^), next get the reduced moment conditions 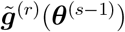 and 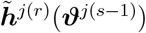, and finally obtain the transformed weighting matrices 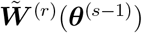 and 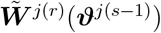.

At the *s*-th iteration, we solve

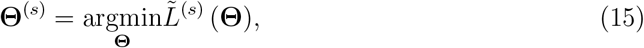

where 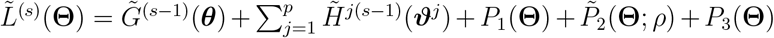. The quadratic form 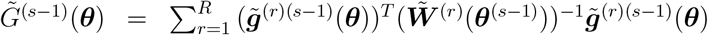 with 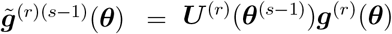 uses principal component matrix ***U*** ^(*r*)^(***θ***^(*s*−1)^) and transformed weighting matrix 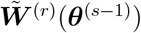 obtained at the (*s* − 1)-th iteration. The quadratic form 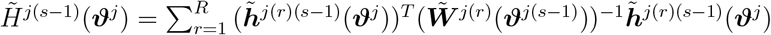 uses principal component matrix ***U*** ^*j*(*r*)^(***ϑ***^*j*(*s*−1)^) and transformed weighting matrix 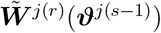 obtained at the (*s* − 1)-th iteration. The above calculation process can be summarized in Algorithm 1, and an expanded version of this algorithm is shown in the Supplementary Material.

The turning parameter *a* in (9) is set to be 3.7 following Fan and Li (2001) and *b* in (9) is set to be 10 following Xue et al. (2022). To determine the regularization parameters ***λ*** = (*λ*_1_, *λ*_2_, *λ*_3_)^*T*^, we propose the following Bayesian-type information criterion (Xue and Qu 2021, Xue et al. 2022):

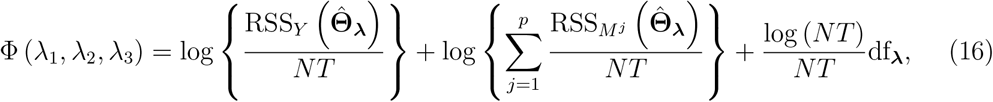

#### Algorithm 1

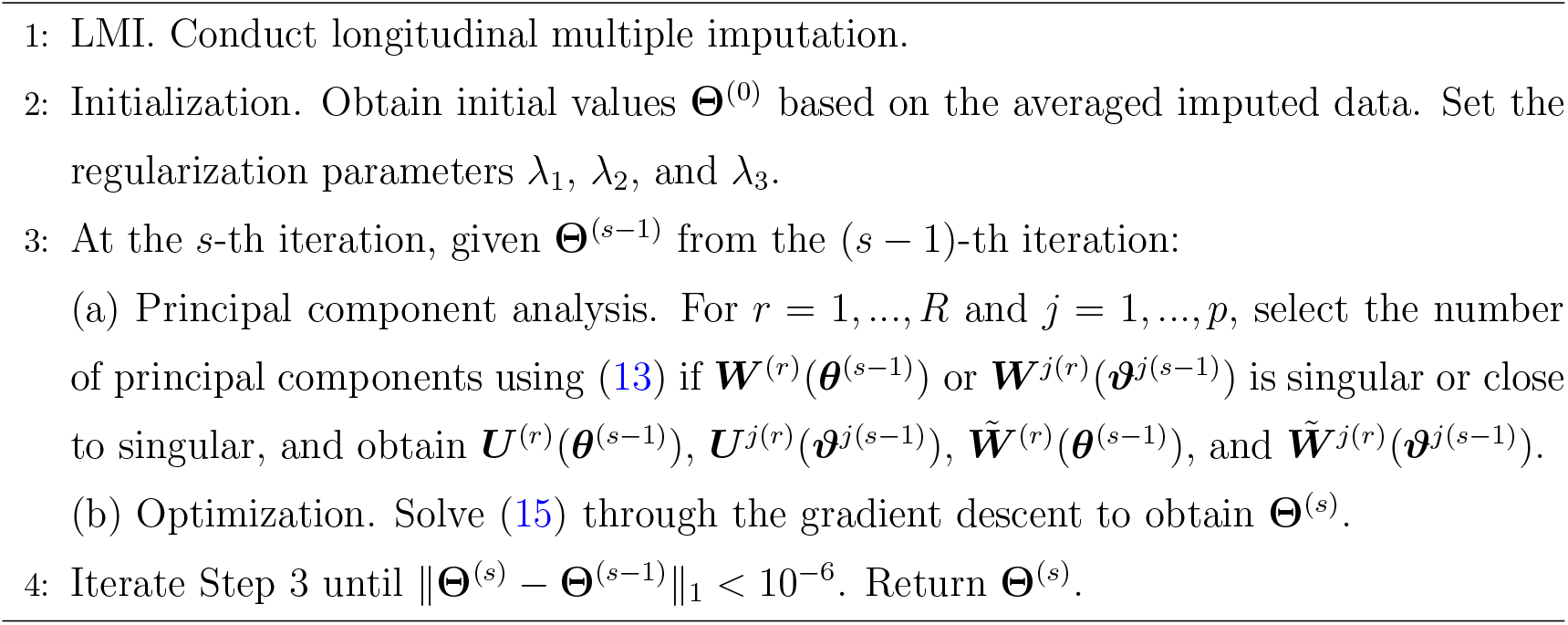

where 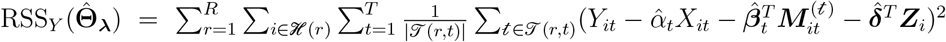 and 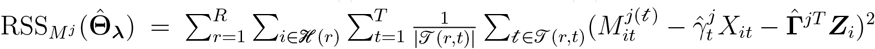 are the residual sum of squares based on imputed data corresponding to models (1) and (2). Degrees of freedom 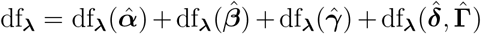, where 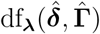 is the number of nonzero estimated values in 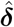 and 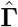, and

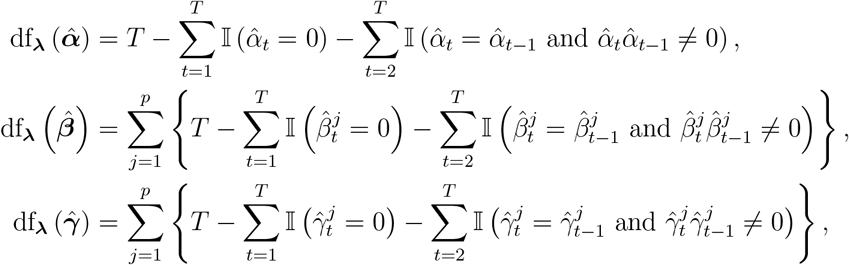

are numbers of nonzero and time-varying estimated values in 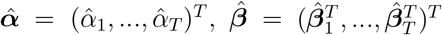, and 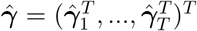, respectively.

### Remark 3.

There are three regularization parameters (*λ*_1_, *λ*_2_, *λ*_3_) that need to be selected during computation, and three-dimensional searches may be computationally costly. In practical applications, we can adjust for confounders ***Z***_*i*_ in models (1) and (2) during the data preprocessing stage (Schuurmans et al. 2024), and then use the residuals for subsequent analyses. This approach removes the need to include parameters ***δ*** and **Γ** in the model, thereby eliminating the necessity of selecting *λ*_3_. For the remaining parameters (*λ*_1_, *λ*_2_), we perform a two-dimensional grid search to identify the combinations which minimize the criterion (16).

## 6 Simulations

In this section, we conduct comprehensive simulation studies to compare the performance of proposed method with some existing approaches. We consider the following time-varying structural equation models:

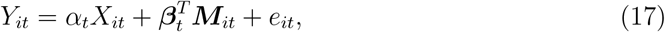

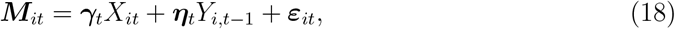

with *q* = 0 and *T* = 4. We generate the exposure *X*_*i*1_ ~ Uniform(0, 1) and *X*_*it*_ = *X*_*i,t*−1_ + Uniform(0, 1) for *t* = 2, …, *T*. The random error vector (*e*_*i*1_, …, *e*_*iT*_)^*T*^ is generated from the multivariate normal distribution 𝒩 (**0, Σ**(*ρ*_1_)), where **Σ**(*ρ*_1_) is the exchangeable correlation structure with parameter *ρ*_1_. The random error vector 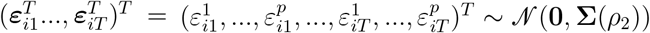, where **Σ**(*ρ*_2_) is the (block-wise) compound symmetry correlation structure with parameter *ρ*_2_. We set *ρ*_1_ = 0.1, *ρ*_2_ = 0.5 and ***η***_*t*_ = **0**.

We consider the following settings with different sample sizes, different dimensions of mediators, different non-monotone missingness structures, and different missingness mechanisms:

**Setting I**. We consider the missingness structure (A) illustrated in Figure 4. We set *N* = 350, *p* = 50, and the sample sizes in group 1-4 are (50, 100, 100, 100). The true parameters 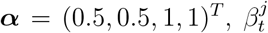 and 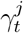 for *j* = 1, …, 14 and *t* = 1, …, 4 are shown in Figure 5. The 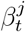 and 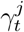 are both zero for *j* = 15, …, 50 and all *t*.

**Figure 4:**
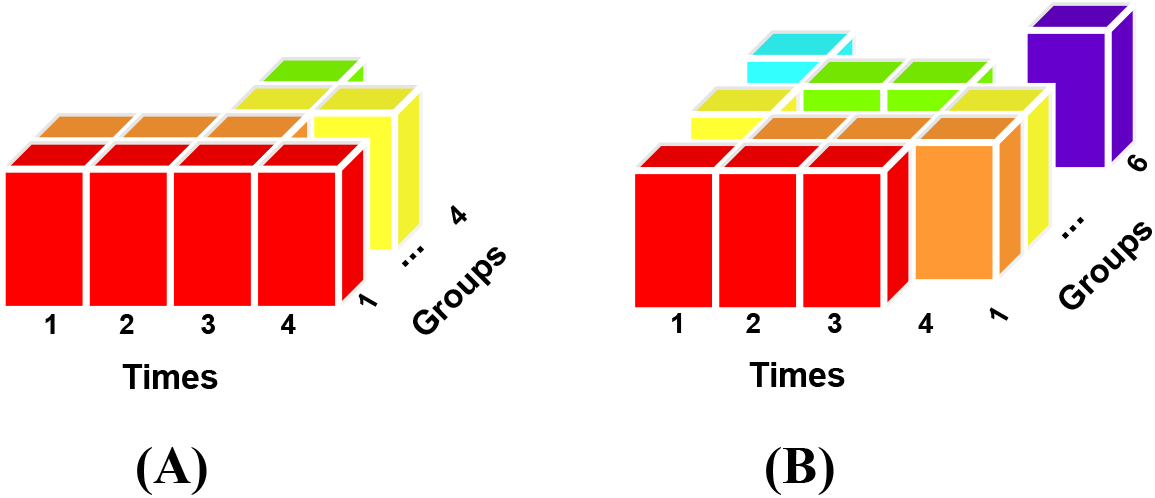
Missingness structures in simulations. Each blank column represents unobserved mediators, while the colored ones represent observed mediators.

**Figure 5:**
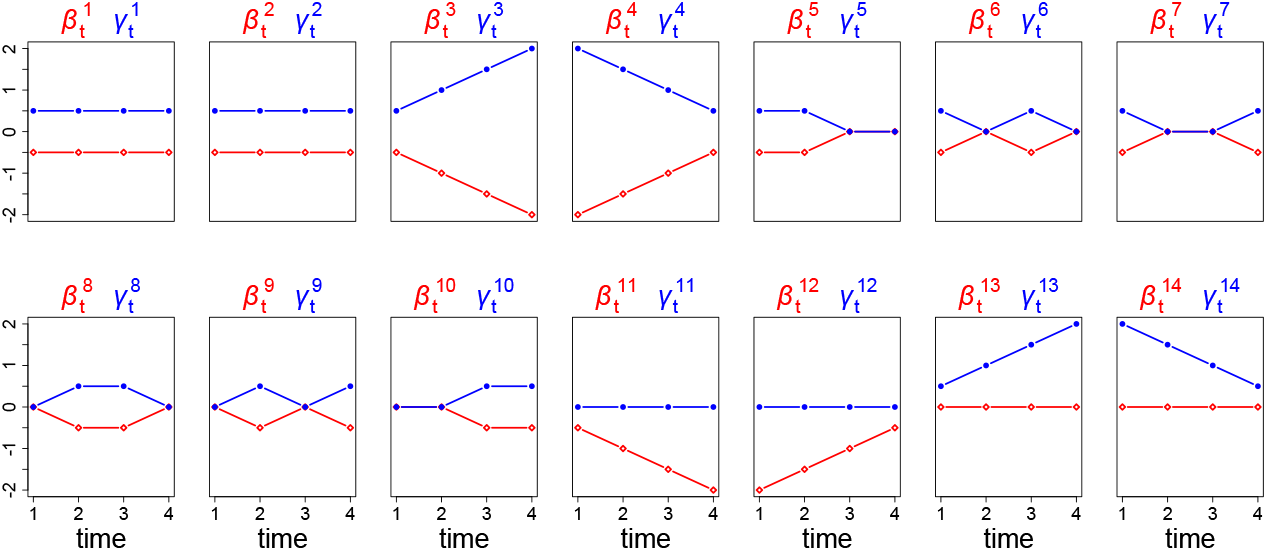
True parameters of 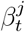 (red colored) and 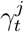 (blue colored) for *j* = 1, …, 14 and *t* = 1, …, 4 in simulation settings I and II. The 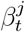 and 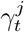 are both zero for *j* = 15, …, 50 and all *t*.

**Setting II**. We consider the missingness structure (B) illustrated in Figure 4, where there is no complete case group. We set *N* = 375, *p* = 50, and the sample sizes in group 1-6 are (75, 75, 75, 50, 50, 50). The true parameters 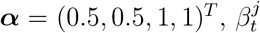 and 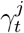 for *j* = 1, …, 14 and *t* = 1, …, 4 are shown in Figure 5. The 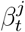 and 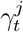 are both zero for *j* = 15, …, 50 and all *t*.

**Setting III**. We consider the missingness structure (A) illustrated in Figure 4. We set *N* = 550, *p* = 200, and the sample sizes in group 1-4 are (100, 150, 150, 150). The true parameters 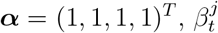 and 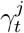 for *j* = 1, …, 22 and *t* = 1, …, 4 are shown in Figure 6. The 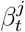 and 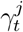 are both zero for *j* = 23, …, 200 and all *t*.

**Figure 6:**
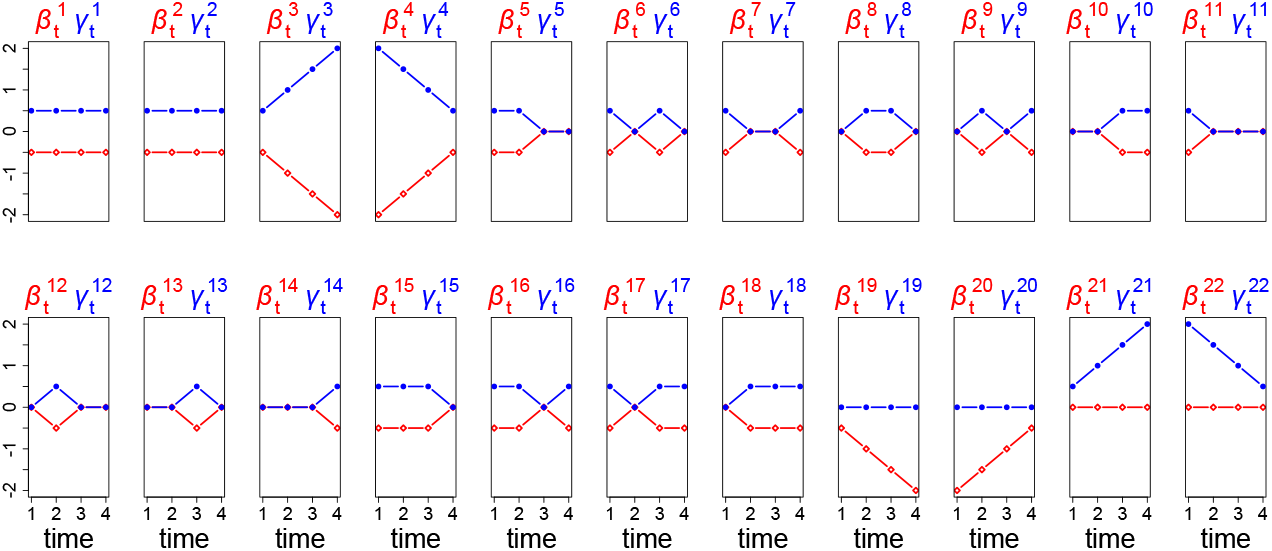
True parameters of 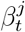 (red colored) and 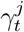 (blue colored) for *j* = 1, …, 22 and *t* = 1, …, 4 in simulation settings III and IV. The 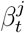 and 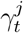 are both zero for *j* = 23, …, 200 and all *t*.

**Setting IV**. We consider the missingness structure (B) illustrated in Figure 4, where there is no complete case group. We set *N* = 675, *p* = 200, and the sample sizes in group 1-6 are (125, 125, 125, 100, 100, 100). The true parameters 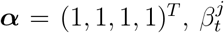 and 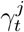 for *j* = 1, …, 22 and *t* = 1, …, 4 are shown in Figure 6. The 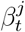 and 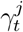 are both zero for *j* = 23, …, 200 and all *t*.

In each of the settings I-IV, we consider three different missingness mechanisms (Little and Rubin 2019): missing completely at random (MCAR), missing at random (MAR), and missing not at random (MNAR). In the MCAR scenario, all subjects are completely randomly assigned to the groups. In the MAR scenario, subjects *i* = 1, …, *N* are sequentially randomly assigned to group 1 with probability proportional to 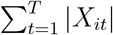, The remaining subjects *i* ∈*/* ℋ(1) are sequentially randomly assigned to group 2 with probability proportional to 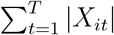, and so on until all subjects are grouped. In the MNAR scenario, the grouping process is the same as that in the MAR scenario, except that the probability is proportional to 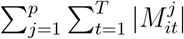.

Our method, using varying-coefficient (VC) models (17) and (18) with LMI-data, is compared with other mediator selection approaches, including: (i) Mix: mixed-effects models with time interactions and lasso penalty (Rijnhart et al. 2022); (ii) Path: linear structural equation models with pathway lasso penalty (Zhao and Luo 2022); (iii) HIMA: linear structural equation models with de-biased lasso and false discovery rate control (Perera et al. 2022); (iv) Bayes: Bayesian sparse models with continuous shrinkage (Song et al. 2020). Note that for VC and Mix, the parameters are estimated using data from all time points simultaneously, while for Path, HIMA, and Bayes, the parameters are estimated using data from each time point separately. We also consider different approaches for handling missing data, including: (i) CC: complete-case analysis which retains only subjects in group 1 with fully observed repeated measurements; (ii) MC: matrix completion via iterative soft-thresholded singular value decomposition (Mazumder, Hastie and Tibshirani 2010); (iii) SI: single imputation based on trajectory means (Jahangiri et al. 2023).

In each replication, to evaluate the accuracy of the mediator selection for each method, we calculate the false negative rate (FNR) 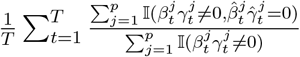 and the false positive rate (FPR) 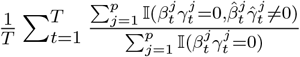, across all time points, where 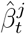 and 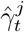 are estimated values of 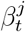 and 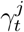 for *t* = 1, …, *T* and *j* = 1, …, *p*. We also calculate the mean squared error (MSE) 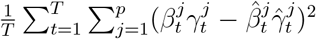 to evaluate the precision of the mediation effect estimation for each method. We say that a method has better performance if it has a smaller FNR+FPR and a smaller MSE.

Tables 1 and 2 summarize the averaged estimation results across 50 replications of different methods for different missingness mechanisms under settings I and II. Tables 1 shows that the proposed VC-LMI method has the smallest FNR+FPR and MSE among all missingness mechanisms, which suggests that the proposed VC-LMI method has the highest accuracy of mediator selection and precision of mediation effect estimation, and its performance is robust to different missingness mechanisms. The VC-CC has a relatively large FNR+FPR, due to the fact that only 50/350 of the subjects with complete repeated measurements are used for analysis. The VC-MC has large FNR+FPR and MSE, possibly because the MC assumes the low-rank structure and noisy entries of the variable matrix, which may not be suitable for our longitudinal data. The VC-SI also has large FNR+FPR and MSE, possibly because the SI utilizes the trajectory mean for imputation, which erases some of the longitudinal changes in variables. Our VC-LMI method makes full use of the correlation among repeated measurements for imputation, and effectively integrates the multiple imputations, which therefore exhibits powerful performance compared with candidate approaches.

**Table 1:** False negative rate (FNR), false positive rate (FPR), FNR+FPR, and mean squared error (MSE) of different methods for different missingness mechanisms under setting I.

|  | missing completely at random |  |  |  | missing at random |  |  |  | missing not at random |  |  |  |
| --- | --- | --- | --- | --- | --- | --- | --- | --- | --- | --- | --- | --- |
|  | FNR | FPR | FNR+FPR | MSE | FNR | FPR | FNR+FPR | MSE | FNR | FPR | FNR+FPR | MSE |
| **VC-LMI** | 0.059 | 0.086 | **0.145** | **0.719** | 0.049 | 0.090 | **0.139** | **0.774** | 0.064 | 0.088 | **0.152** | **0.715** |
| VC-CC | 0.174 | 0.054 | 0.228 | 1.032 | 0.180 | 0.057 | 0.237 | 1.018 | 0.150 | 0.058 | 0.208 | 0.860 |
| VC-MC | 0.419 | 0.072 | 0.491 | 8.332 | 0.333 | 0.074 | 0.407 | 8.590 | 0.401 | 0.071 | 0.471 | 8.176 |
| VC-SI | 0.203 | 0.036 | 0.239 | 5.065 | 0.180 | 0.034 | 0.214 | 5.228 | 0.196 | 0.036 | 0.232 | 5.208 |
| Mix-CC | 0.111 | 0.105 | 0.215 | 2.638 | 0.109 | 0.114 | 0.222 | 2.525 | 0.116 | 0.099 | 0.215 | 2.319 |
| Mix-MC | 0.086 | 0.327 | 0.414 | 8.815 | 0.077 | 0.320 | 0.397 | 9.214 | 0.092 | 0.314 | 0.407 | 8.772 |
| Mix-SI | 0.057 | 0.167 | 0.224 | 5.839 | 0.043 | 0.163 | 0.205 | 6.016 | 0.049 | 0.157 | 0.206 | 5.965 |
| Path-CC | 0.272 | 0.065 | 0.337 | 7.215 | 0.288 | 0.064 | 0.351 | 7.125 | 0.258 | 0.065 | 0.323 | 7.082 |
| Path-MC | 0.560 | 0.032 | 0.592 | 10.420 | 0.470 | 0.028 | 0.498 | 10.364 | 0.541 | 0.033 | 0.574 | 10.382 |
| Path-SI | 0.248 | 0.020 | 0.268 | 9.198 | 0.227 | 0.022 | 0.250 | 9.319 | 0.249 | 0.021 | 0.270 | 9.285 |
| HIMA-CC | 0.679 | 0.005 | 0.685 | 1.715 | 0.702 | 0.004 | 0.707 | 2.198 | 0.669 | 0.004 | 0.673 | 1.708 |
| HIMA-MC | 0.574 | 0.035 | 0.609 | 8.742 | 0.489 | 0.035 | 0.524 | 9.173 | 0.547 | 0.040 | 0.587 | 8.579 |
| HIMA-SI | 0.376 | 0.014 | 0.390 | 4.895 | 0.361 | 0.015 | 0.377 | 5.066 | 0.370 | 0.015 | 0.385 | 5.158 |
| Bayes-CC | 0.916 | 0.000 | 0.916 | 5.577 | 0.916 | 0.000 | 0.916 | 5.508 | 0.901 | 0.000 | 0.901 | 4.797 |
| Bayes-MC | 0.786 | 0.019 | 0.805 | 10.043 | 0.754 | 0.021 | 0.775 | 10.840 | 0.760 | 0.020 | 0.780 | 9.711 |
| Bayes-SI | 0.689 | 0.001 | 0.690 | 6.129 | 0.675 | 0.001 | 0.676 | 6.129 | 0.659 | 0.001 | 0.660 | 6.115 |

Note: Each estimator is denoted as “mediator selection method - missing data handling method”. Results are averaged across all time points and 50 replications.

**Table 2:** False negative rate (FNR), false positive rate (FPR), FNR+FPR, and mean squared error (MSE) of different methods for different missingness mechanisms under setting II.

|  | missing completely at random |  |  |  | missing at random |  |  |  | missing not at random |  |  |  |
| --- | --- | --- | --- | --- | --- | --- | --- | --- | --- | --- | --- | --- |
|  | FNR | FPR | FNR+FPR | MSE | FNR | FPR | FNR+FPR | MSE | FNR | FPR | FNR+FPR | MSE |
| **VC-LMI** | 0.046 | 0.105 | **0.151** | **0.790** | 0.048 | 0.113 | **0.160** | **0.855** | 0.056 | 0.107 | **0.163** | **0.755** |
| VC-MC | 0.439 | 0.076 | 0.514 | 9.195 | 0.369 | 0.070 | 0.439 | 9.082 | 0.426 | 0.072 | 0.498 | 8.962 |
| VC-SI | 0.254 | 0.043 | 0.297 | 5.426 | 0.276 | 0.043 | 0.319 | 5.477 | 0.246 | 0.043 | 0.290 | 5.374 |
| Mix-MC | 0.105 | 0.330 | 0.435 | 10.002 | 0.091 | 0.291 | 0.382 | 10.131 | 0.114 | 0.297 | 0.411 | 9.880 |
| Mix-SI | 0.057 | 0.140 | 0.198 | 6.707 | 0.060 | 0.143 | 0.203 | 6.606 | 0.056 | 0.151 | 0.207 | 6.581 |
| Path-MC | 0.646 | 0.022 | 0.668 | 10.908 | 0.559 | 0.017 | 0.577 | 10.770 | 0.611 | 0.018 | 0.630 | 10.851 |
| Path-SI | 0.286 | 0.019 | 0.305 | 9.609 | 0.276 | 0.016 | 0.292 | 9.534 | 0.264 | 0.019 | 0.283 | 9.544 |
| HIMA-MC | 0.654 | 0.045 | 0.698 | 9.667 | 0.571 | 0.046 | 0.618 | 9.837 | 0.631 | 0.048 | 0.680 | 9.564 |
| HIMA-SI | 0.457 | 0.021 | 0.478 | 5.321 | 0.461 | 0.018 | 0.479 | 5.287 | 0.429 | 0.021 | 0.450 | 5.360 |
| Bayes-MC | 0.915 | 0.028 | 0.943 | 10.791 | 0.876 | 0.031 | 0.907 | 11.405 | 0.902 | 0.030 | 0.933 | 10.800 |
| Bayes-SI | 0.756 | 0.008 | 0.764 | 5.960 | 0.774 | 0.005 | 0.780 | 6.075 | 0.764 | 0.006 | 0.770 | 6.010 |

Note: Each estimator is denoted as “mediator selection method - missing data handling method”. Results are averaged across all time points and 50 replications.

For other mediator selection methods in Tables 1, both FNR+FPR and MSE are large. Specifically, the Mix shows large FPR, while the Path provides large FNR. The reason for their poor estimations may be because the mediator selection methods with some shortcomings (too many interactions in Mix and over-shrinkage in Path) are used on the datasets with poor quality. The HIMA and Bayes provide a smaller FPR but a larger FNR, possibly due to the multiple-testing procedure for HIMA and the posterior inclusion probability for Bayes being overly conservative.

Table 2 summarizes the estimation results under setting II corresponding to the missingness structure (B) in Figure 4. Since no subject has complete repeated measurements, the CC is not applicable. Our proposed VC-LMI method has the smallest FNR+FPR and MSE among all candidate approaches. The estimation results under settings III and IV in the high-dimensional context are provided in Tables S1 and S2 in the Supplementary Material, which show that our proposed VC-LMI method also beats other candidate approaches, based on its smallest FNR+FPR and MSE. The Supplementary Material includes additional simulations, where Tables S3 and S4 demonstrate that our method continues to outperform other candidate approaches in higher-dimensional settings (*p* = 2000) and scenarios with more time points (*T* = 5), respectively. Table S5 shows that combining multiple imputations through the GMM outperforms approach that selecting imputations with the smallest variance.

## 7 DNHS data analysis

In this section, we investigate how DNAm dynamically mediates the effect of traumatic experiences on development of PTSD, utilizing the DNHS data. Specifically, we treat the number of traumatic events as exposure *X*_*it*_, the logarithm transformed PTS symptom severity score as outcome *Y*_*it*_, and the M-value transformed (Kruppa et al. 2021) DNAm CpG sites as potential mediators ***M***_*it*_. The study includes 526 subjects in waves 1, 2, 4, and 5, which are referred to as time points 1-4. In our data, exposure *X*_*it*_ and outcome *Y*_*it*_ are always observed, but the mediators ***M***_*it*_ suffer from non-monotone missingness as shown in Figure 1: ***M***_*i*_ of some subjects may be available at certain visits, but missing at the next time point, and measured again at later visits.

There are two steps in the data preprocessing phase. First, we adjust some demographic information and blood work measured at baseline. For the outcome corresponding to model (1), we adjust age, gender, race, education, income, cigarette use, depression, and social cohesion score (Johns et al. 2012, Wani et al. 2021). For the mediators corresponding to model (2), we adjust age, gender, cigarette use, CD4 T cells, CD8 T cells, natural killer cells, B cells, and monocytes (Occean et al. 2022). The residuals after adjustment and the models (17) and (18) are used for subsequent analyses. Second, given the limited number of subjects and the ultra-high dimensional potential mediators (1879 DNAm CpG sites), we carry out a screening process to reduce the dimension to a moderate scale below the sample size (Fan and Lv 2008). Specifically, for *t* = 1, …, *T* and *j* = 1, …, *p*, we consider a series of marginal models 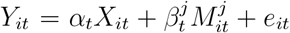 and 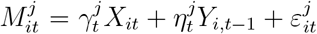. Along the lines of the sure independence mediator screening (Perera et al. 2022), for each time point *t* = 1, …, *T*, we use the subjects with observed mediators to fit the marginal models, then obtain a subset 𝒟_*t*_ ={*j*: *M*^*j*^ is among the top [*n*_*t*_*/*log(*n*_*t*_)] largest 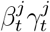 effect}, where *n*_*t*_ is the sample size at time point *t* and [·] denotes the integer part. Then we take 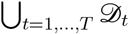 as the index set of potential mediators after screening. There are 153 DNAm CpG sites remaining for the next step of analysis.

We use the proposed VC-LMI method to fit the preprocessed data. The estimated direct effect is 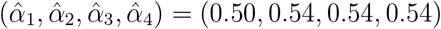, which means that trauma exposure has a sustained direct effect on PTSD. Figures 7 and 8 show the 13 selected CpG sites, each of which has nonzero 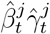 at least at one time point. We can find that certain CpG sites (e.g., cg22564046) initially have nonzero mediation effects, but the effects disappear over time. Certain CpG sites (e.g., cg17318247) do not show mediation effects in the early stages, but the effects emerge over time. This indicates that the effects at the CpG level may vary in different stages of the disease process.

**Figure 7:**
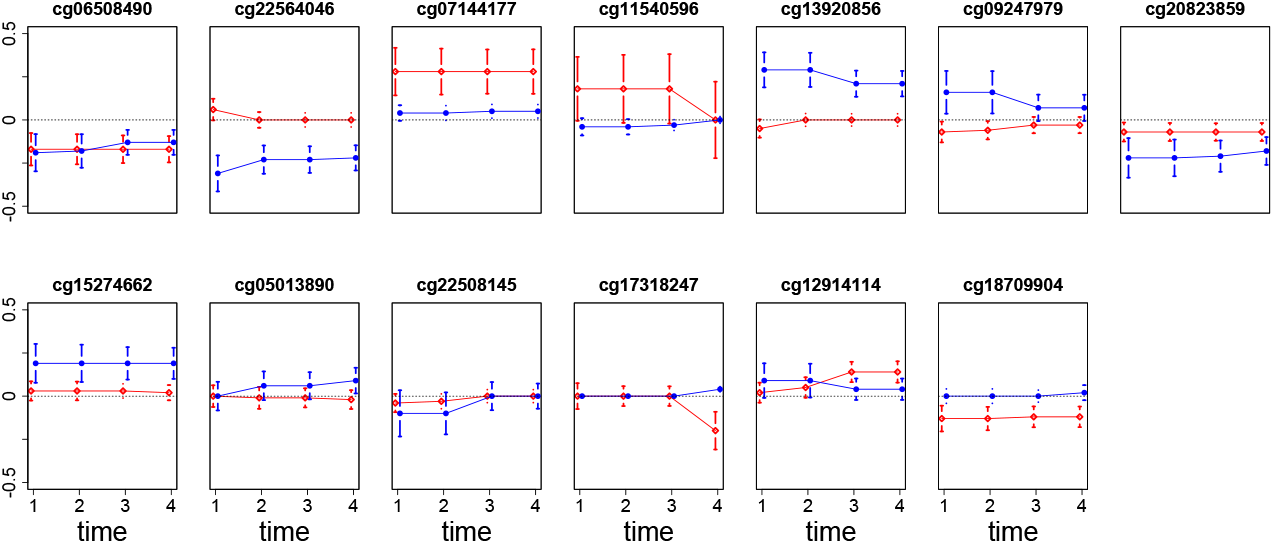
Estimated values 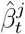 (red colored) and 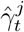 (blue colored) with Bootstrap 95% confidence intervals for the selected CpG sites and *t* = 1, …, 4, based on the proposed VC-LMI method.

**Figure 8:**
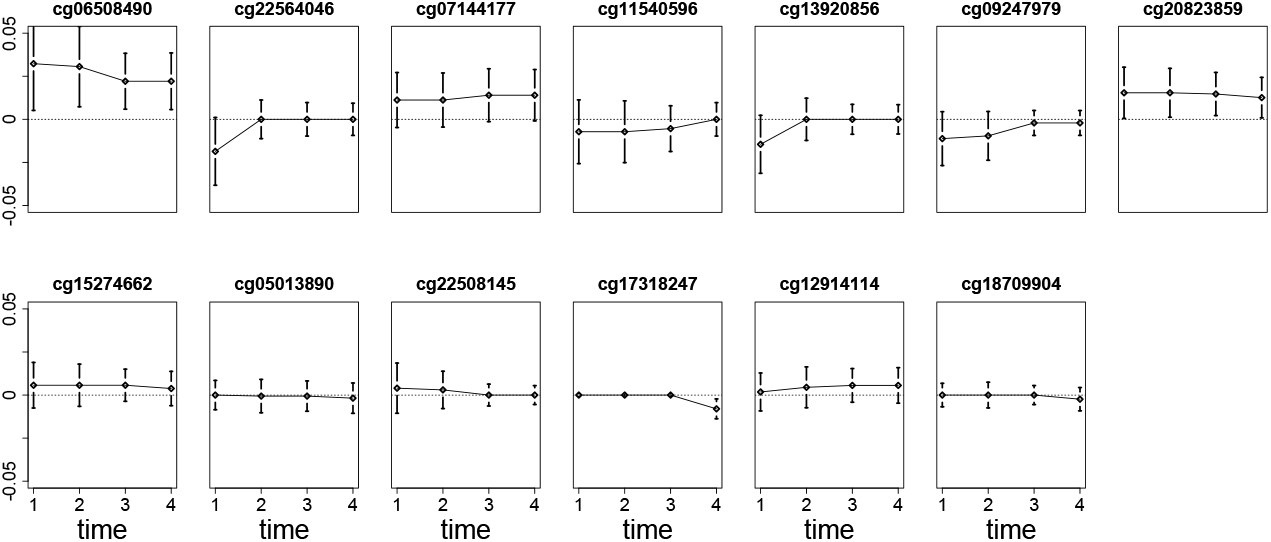
Estimated mediation effects 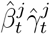 with Bootstrap 95% confidence intervals for the selected CpG sites and *t* = 1, …, 4, based on the proposed VC-LMI method.

As shown in Figure 8, the estimated mediation effects for certain CpG sites at specific time points, such as cg22508145 at *t* = 1, 2 and cg09247979 at *t* = 3, 4, appear small or close to zero. These seemingly negligible effects should be interpreted within the broader context of longitudinal mediation analysis and the cumulative role of DNAm across multiple time points. First, mediation effects can vary over time due to the dynamic nature of DNAm and its interactions with environmental exposures and disease progression. Small mediation effects at individual time points do not necessarily indicate that the CpG site lacks relevance as a mediator. Instead, they may reflect temporal fluctuations or periods during which the indirect effect is less pronounced. Second, while mediation effects may appear negligible at specific time points, the cumulative contribution of a CpG site across all time points could still be meaningful. Longitudinal analyses are designed to capture these aggregate patterns, which may not be evident when focusing solely on isolated single-time-point effects. Finally, even small mediation effects can hold biological significance, especially in high-dimensional settings where individual mediators contribute subtly but collectively to disease mechanisms (Xue et al. 2022).

Table 3 lists the gene information corresponding to the selected CpG sites. Eleven of these sites are annotated to human genes according to the Illumina annotation files (*CAT, PFKP, EEF1E1, TANC1, PTPRK, OVGP1, RPS6KA2, CPAMD8, CDK16, FAM120B*, and *C14orf182*). Of these eleven genes, *PTPRK* has been shown to be associated with mental health disorders such as PTSD (Chitrala, Nagarkatti and Nagarkatti 2016), *RPS6KA2* has been reported as significant KEGG pathways for PTSD in epigenome-wide DNA methylation study (Kuan et al. 2017). Other CpG sites while not located near any protein-coding genes, fall within ENCODE candidate Cis-Regulatory Elements that are adjacent to highly conserved non-coding genomic regions (Moore et al. 2020).

**Table 3:** Gene information corresponding to the selected CpG sites.

| CpG | chromosome | Gene |
| --- | --- | --- |
| cg06508490 | 11 | CAT |
| cg22564046 | 10 | PFKP |
| cg07144177 | 6 | EEF1E1 |
| cg11540596 | 6 | NA/Desert |
| cg13920856 | 2 | TANC1 |
| cg09247979 | 6 | PTPRK |
| cg20823859 | 1 | OVGP1 |
| cg15274662 | 7 | NA/Desert |
| cg05013890 | 6 | RPS6KA2 |
| cg22508145 | 19 | CPAMD8 |
| cg17318247 | X | CDK16 |
| cg12914114 | 6 | FAM120B |
| cg18709904 | 14 | C14orf182 |

Beyond the time-invariant associations, our results further show that the effects of genes change over time, which may reveal how the biological embedding of trauma exposure through DNAm contributes to PTSD risk over development. Our findings could have important clinical implications for time-sensitive risk prediction in trauma-exposed populations.

To compare the performance of the proposed VC-LMI method with other approaches, we randomly split the preprocessed data into a training set (80% of subjects) and a testing set (20% of subjects; the index set is denoted as T) for 50 replications. In each replication, for each method, we train the model on the training set and obtain the estimated values of parameters. Then we calculate the mean number of selected mediators (No.-mediator) as 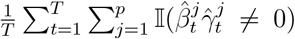, the prediction mean squared error (PMSE-*β*) for the outcome model (17) as 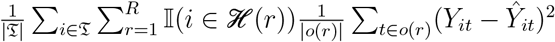, and the prediction mean squared error (PMSE-*γ*) for the mediator model (18) as 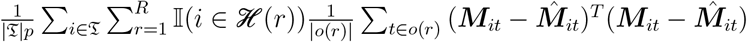, on the testing set across all time points, where 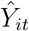 and 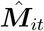 are the fitted values of *Y*_*it*_ and ***M***_*it*_ for *i* ∈ ℐ and *t* ∈ *o*(*r*). Table 4 summarizes the averaged estimation results across 50 replications. We observe that our VC-LMI method has the smallest PMSE both in the outcome model and the mediator model, indicating that the proposed method is more accurate in terms of prediction. For the Mix and the Path, despite more mediators being selected based on the imputed dataset, the PMSE is larger, possibly due to overfitting. For the HIMA and the Bayesian approaches, no mediator passes their overly conservative significance tests, which is consistent with the simulation results in Section 6.

**Table 4:**
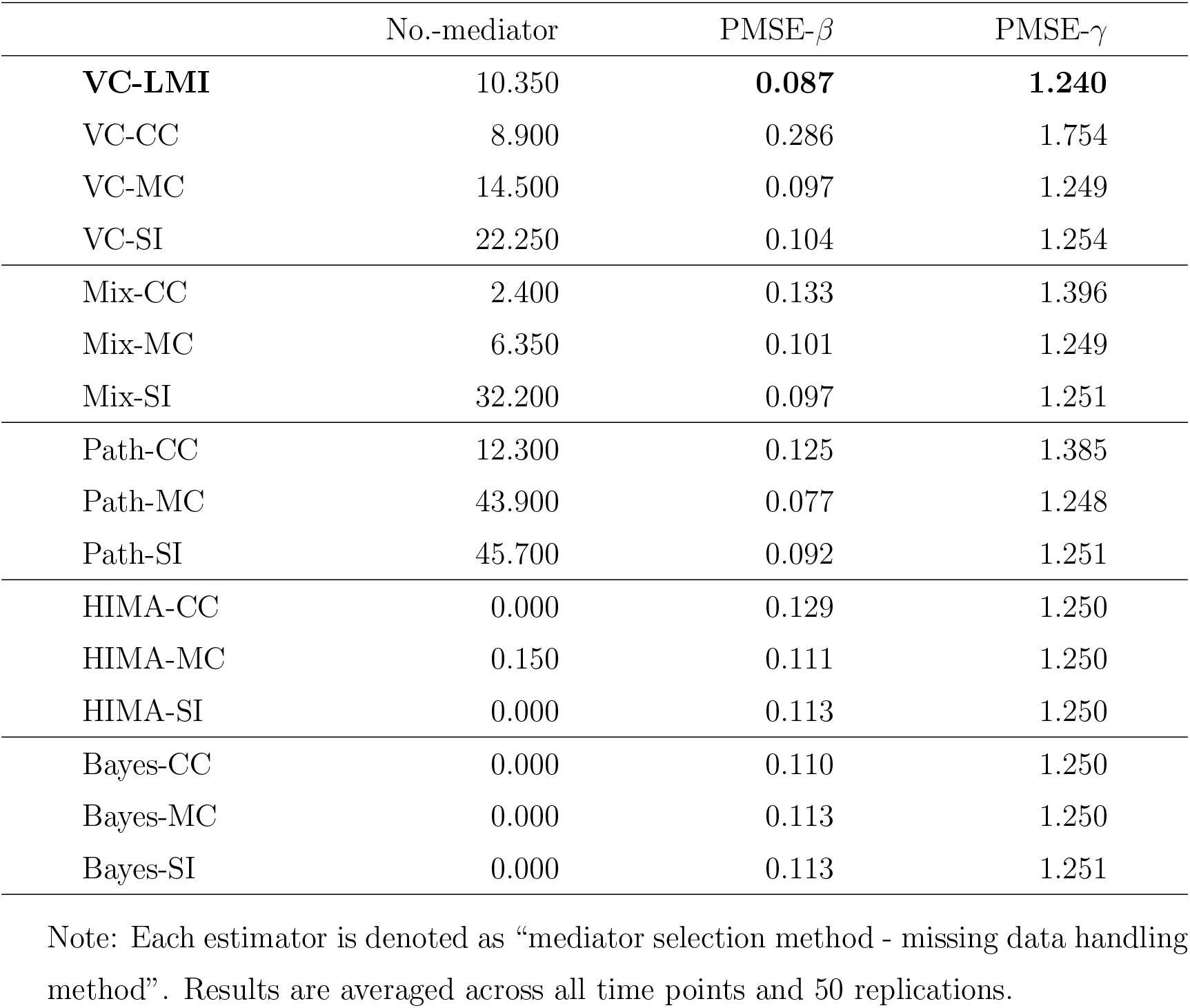
Mean number of selected mediators (No.-mediator), prediction mean squared error (PMSE-*β*) for the outcome model (17), and prediction mean squared error (PMSE-*γ*) for the mediator model (18) in DNHS data analysis.

## 8 Discussion

In this paper, to investigate the time-varying mediation effects of DNAm on the relationship between trauma and PTSD, we propose time-varying structural equation models. The LMI approach is proposed to handle non-monotone missing DNAm and the regularization approach is incorporated to select relevant mediators and capture time-varying effects. In simulations, the proposed method compares favorably against existing competitors in various scenarios. In DNHS data analysis, we successfully identify potential DNAm CpG sites which show dynamic mediation effects.

We assume the mediators 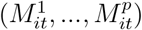 are correlated but we do not specify causal ordering. The main reason for this assumption is that, based on the existing literature, external adverse environmental exposures or disease conditions are likely to influence DNAm across multiple CpG sites (Turecki and Meaney 2016), whereas the relationships between DNAm levels across CpG sites are primarily correlational (Mou et al. 2022). Such assumptions of parallel relationships among mediators are commonly made in high-dimensional DNAm-related mediation analysis (e.g., Song et al. 2020, Perera et al. 2022, Xue et al. 2022). Furthermore, even if causal ordering existed among high-dimensional DNAm CpG sites, establishing the correct causal order is infeasible in complex biological systems, instead they are characterized by intricate interactions and feedback loops (Tai et al. 2022). In our study, we also lack sufficient genetic knowledge to reliably support causal ordering. A more refined understanding of the causal relationships among high-dimensional mediators may require the application of directed acyclic graph learning techniques (Shi and Li 2022), which might be able to identify the complex causal structures among variables. However, such analyses extend beyond the scope of this study and could be explored in future research.

Unmeasured confounding is an inherent challenge in population-based studies. In our DNHS data analysis, we incorporate a comprehensive set of demographic information and blood work measured at baseline to control for confounding bias as much as possible. Nonetheless, some unmeasured confounders, such as participants’ other medical treatments or psychological interventions for PTSD not captured in the DNHS investigation, may still exist. To address these remaining unmeasured confounders, methods such as instrumental variable approaches (Chen et al. 2023), proxy causal learning frameworks (Dukes et al. 2023), or de-confounding techniques (Yuan and Qu 2024) could be employed. Exploring these methodologies could be a promising direction for future research.

In our imputation step, we estimate 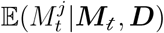 using specific groups that include observed 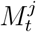 and ***M***_*t*_, then impute the unobserved 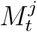 in other groups that contain observed ***M***_*t*_ but missing 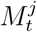. The rationale for our imputation method relies on the premise that the conditional distributions of mediators in the observed data are the same as those in the missing data, which holds only under the assumptions of MCAR or MAR. For mediators that are MNAR, the direct and indirect effects corresponding to the coefficients in models (1) and (2) are not identifiable without additional assumptions. Potential strategies include modeling the missing indicator 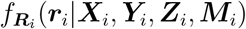 based on the completeness assumption (Zuo et al. 2024), or introducing instrumental or shadow variables to impose specific structures on the missingness mechanism (Shan et al. 2024). Extending our method to accommodate MNAR mediators using these strategies, as well as further addressing missing outcomes (Qin et al. 2019), are important directions for future research.

For the statistical inference of the mediation effect, there are two main challenges including the limiting distributions of the estimators for parameters in models (1) and (2), and the composite *p*-value of the mediation effect. Specifically, the penalty functions in (9) and (10) may present challenges in deriving the asymptotic distribution. Some existing studies used the bi-level penalty similar to that of *P*_1_(**Θ**) in (9) and the corresponding theoretical results show that the estimator of nonzero coefficients in general converges to nonnormal distribution (Huang et al. 2009). Our estimator includes an additional fusion penalty *P*_2_(**Θ**) in (10) to serve the purpose of shrinking similar effects at adjacent time points, which may lead to more complex limiting distributions. On the other hand, to test whether *M*^*j*^ mediates the effect of *X* on *Y*, the null and alternative hypotheses can be formulated as 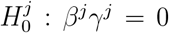 versus 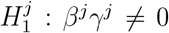. The null hypothesis encompasses a product of parameters, which is composite and consists of three cases as case 1: *β*^*j*^ = 0 and *γ*^*j*^ ≠ 0, case 2: *β*^*j*^ ≠ 0 and *γ*^*j*^ = 0, and case 3: *β*^*j*^ = 0 and *γ*^*j*^ = 0. Constructing the test statistic and deriving the reference distribution and composite *p*-value under the composite null hypothesis are still open problems in the field of high-dimensional mediation analysis, and relevant studies can be found in the review by Du et al. (2023). Considering the complex limiting distributions of 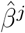 and 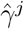, it is more challenge to conduct statistical inference for the mediation effect 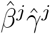. We will develop theories and inference tools of the proposed estimator in our future research.

## Funding

The research is supported by NSF Grant DMS-2210640 and 1952406, and in part by NIH Grant R01MD011728.

## Supplement

### Supplement to “Time-varying mediation analysis for incomplete data with application to DNA methylation study for PTSD”

The Supplementary Material includes three sections. Section 1 contains the GMM estimator corresponding to each mediator *M*^*j*^ (*j* = 1, …, *p*) in model (2). Section 2 contains the expanded version of Algorithm 1. Section 3 contains the additional simulation results.

### Supplemental code

We have implemented the proposed longitudinal multiple imputation method as an R function named “LMI” and the generalized method of moments with principal component analysis as another R function named “GMM PCA”. Simulation code, data, and application code, along with detailed instructions, are provided.

## Notes

### Competing Interest Statement

The authors have declared no competing interest.

### Summary of Updates

The revised manuscript includes the updated model, simulation results, and real data analysis results.

